# Assessing Protein Surface-Based Scoring for Interpreting Genomic Variants

**DOI:** 10.1101/2023.12.22.573111

**Authors:** Nikita R. Dsouza, Neshatul Haque, Swarnendu Tripathi, Michael T. Zimmermann

## Abstract

Clinical genomics sequencing is rapidly expanding the number of variants that need to be functionally elucidated. Interpreting genetic variants (i.e., mutations) usually begins by identifying how they affect protein-coding sequences. Still, the 3D protein molecule is rarely considered for large-scale variant analysis, nor for their effects on how proteins interact with each other and their environment. We propose a standardized approach to scoring protein surface property changes as a new dimension for functionally and mechanistically interpreting genomic variants. Further, it directs hypothesis generation for functional genomics research to learn more about the encoded protein’s function. We developed a novel method leveraging 3D structures and time-dependent simulations to score and statistically evaluate protein surface property changes. We evaluated positive controls composed of 8 thermophilic versus mesophilic orthologs and variants that experimentally change protein solubility, which all showed large and statistically significant differences in charge distribution (p<0.01). We scored static 3D structures and dynamic ensembles for 43 independent variants (23 pathogenic and 20 uninterpreted) across four proteins. Focusing on the potassium ion channel, KCNK9, average local surface potential shifts were 0.41 k_B_T/ec with an average p-value of 1×10^−2^. In contrast, dynamic ensemble shifts averaged 1.15 k_B_T/ec and an average p-value of 1×10^−5^, enabling the identification of changes far from mutated sites. This study demonstrates that an objective assessment of how mutations affect electrostatic distributions of protein surfaces can aid in interpreting genomic variants discovered through clinical genomics sequencing.

## Introduction

Clinical genomics data is increasingly gathered to diagnose diseases and identify potential treatment options. Thus, interpreting genomics data is critical for Precision Medicine ^1-3^. Most genomic variants used in clinical decision-making alter the protein-coding sequence of genes, yet properties of the 3D and time-dynamic molecule are rarely assessed ^4^. Proteins interact with each other and bind to metabolites and drugs using specific surfaces. Enzymatic sites, for example, have precisely calibrated surfaces for their chemistry. Similarly, many interaction surfaces, such as those that form multiprotein complexes, have been tuned to accommodate folding, assembly, and functional motions. Protein surface properties are thus critical characteristics for function.

Currently, interpreting genomic variants considers gene or protein sequence changes, with data such as prior observation of variants, protein domains, or amino acid conservation annotated to those sequences ^5-7^. However, it frequently occurs that the polypeptide chain folds on itself, making a diverse array of loops, turns, twists, and bends, resulting in regions of protein surfaces that are non-linear in the protein sequence. Such characters, with unique charge patterns, hydrophobicity, and shape, become characteristic features of the protein that are indispensable for the protein’s function. These surfaces are distinctive, irregular, and based on their specific role, such as ligand binding, protein-protein interaction, protein-membrane interaction, *etc*. Therefore, a 3D approach is more appropriate for the evaluation of variants.

Surface properties are determined by the combined contributions of intrinsic and extrinsic factors, which influence each other in a dynamic interplay of entropic and enthalpic exchange. In an absolute sense, Anfinsen’s dogma would reduce protein intrinsic factors to the amino acid sequence. Still, the sequence itself can be separated into components of backbone dihedral angles, sidechain chemistry, sidechain packing, folding kinetics, and more. Protein extrinsic factors include substrates, cofactors, ions, other proteins, lipids, carbohydrates, and molecules that make up the environment. Further, surfaces have different characteristics, including shape, flexibility, and how hydrophobic and electrostatic potentials distribute across them. Because protein intrinsic and extrinsic factors constantly interplay, mutations can tip the balance among their combinations, potentially altering or dysregulating function. In general, any small perturbation in intrinsic or extrinsic factors leads to cooperative surface rearrangements ^8^, which could vary depending on the type and nature of the perturbation. While surface differences have been investigated for specific proteins and disease variants ^9^, there are no general and standardized approaches for statistically determining the significance of surface changes for genetic variants identified through high-throughput sequencing. Therefore, this study seeks to determine the statistical performance of a strategy for scoring differences in protein surface electrostatic properties due to mutation.

We designed this study, to begin with straightforward positive controls for global and local changes, then progress to mutations with known effects, followed by genomic variants currently uninterpreted in the same genes. The positive control variants *a priori* should have distinct changes, such as thermophilic and mesophilic orthologs, and experimentally determined mutations that increase protein solubility. Then, equipped with a reference for what functional surface changes can look like, we characterized a series of genomic variants identified using high-throughput sequencing of patients, undiagnosed at the time of sequencing, but through our collaborative functional genomic studies ^10^ likely having damaging variants causing rare genetic diseases, and pathogenic controls in the same proteins. We used computational mutagenesis for each variant and scored their corresponding changes to protein surfaces using static models of each protein. Finally, we used physics-based simulations to model protein dynamic ensembles and compared results using these two levels of detail of the protein surface. Our data demonstrates that greater sensitivity is gained by including protein dynamics. Thus, we conclude that it is possible to standardize a process for scoring genomic mutations using protein surface-based scores, adding a new dimension to the interpretation of human genetic variation.

## Methods

### Aggregating Genomic Variants

We initiated the current study through collaborative work in rare and undiagnosed diseases, where patient cohorts were accrued based on common phenotypes and variants of uncertain significance (VUS) within genes plausible for causing the phenotype. The cohort studied are TBL1XR1^11^, KCNK9^12^, and PIK3R1^13^. To gather additional genomic variants in the same genes, we used BioR^14^ and custom scripts to map them onto 3D protein structures. Variants were obtained from GnomAD^10^, HGMD^15^, ClinVar^16^, and COSMIC^17^.

### Experimental Structures of Wild-Type and Genomic Mutations

We selected four groups for comparison (**Figure 1**). First, thermophilic and mesophilic pairs and solubilizing mutations were used as two types of positive controls to parametrize the statistical tests described below. Then, we selected 3D static structures of disease-associated genomic variants as a third group. Our fourth group consisted of the same set of mutations but with the additional data of protein dynamic ensemble.

**Figure 1:**
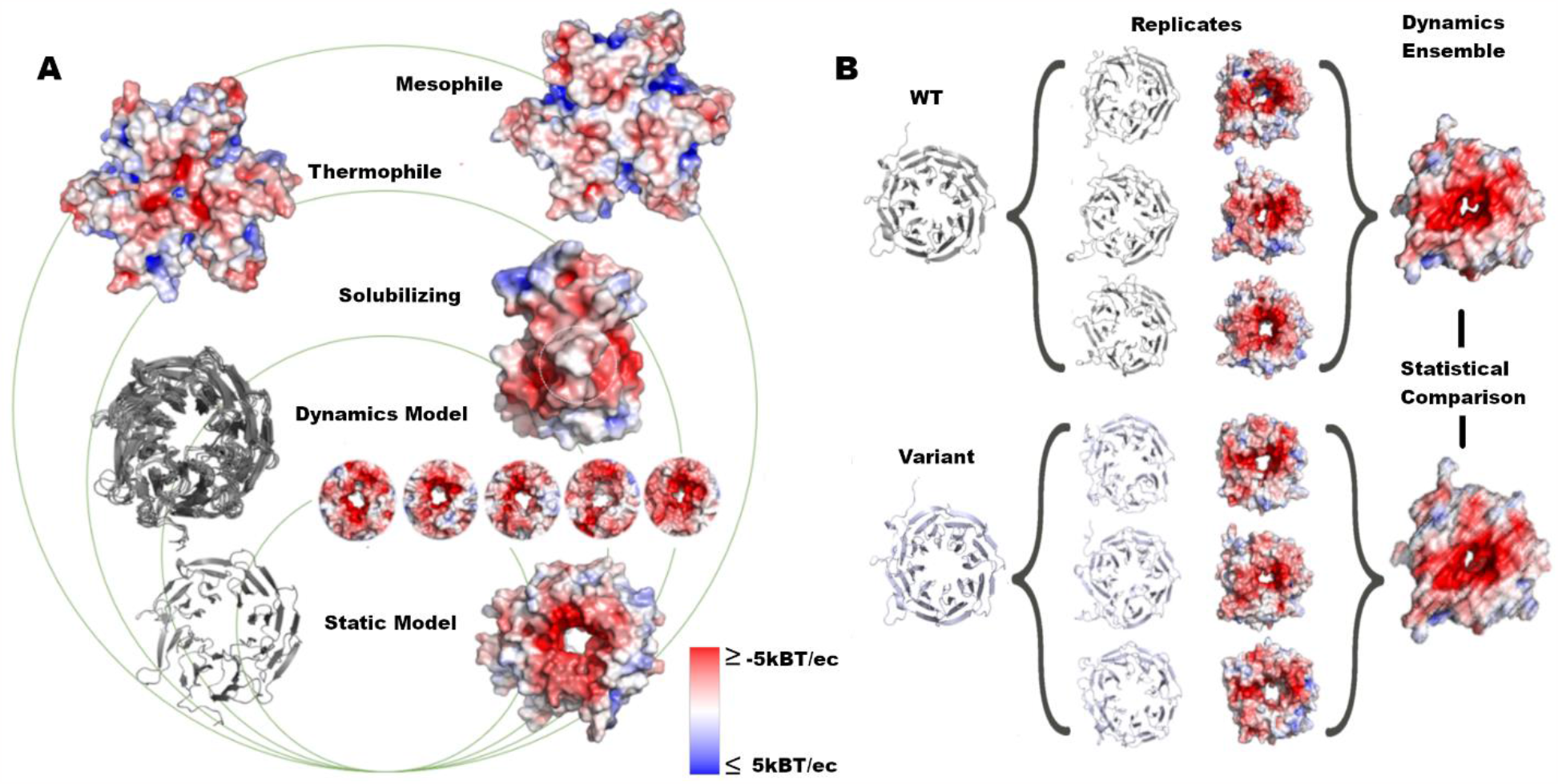
Assessment of structure and dynamics of protein surfaces. **A)** In this work, we analyzed the surface changes of proteins with groups ranging from global to local changes caused by variants. **B)** We ran MD simulations in triplicate, calculated the electrostatic potential for each replicate, and measured the difference in the dynamic ensemble for each structure.

When extant, we used experimental structures for each protein obtained from the Protein Data Bank (PDB)^18^. **Table 1** contains the PDB IDs of the thermophiles and mesophiles used for this study. **Table 2** provides PDB IDs for proteins and genetically engineered point mutations that showed a substantial change in solubility, likely caused by changing surface properties. **Table 3** and **Table 4** have the PDB IDs for the WT structures of the proteins and two genetically engineered mutations, G318R and K297N^19^, in UROD (3gvq). Experimentally solved structures existed for Wild Type (WT) TBL1XR1 (4lg9)^20^ and IGF1R (1m7n^21^), so we used homology-based methods to fill in the loops that were not resolved^22,23^. We used I-Tasser^24^ to generate the model for KCNK9 from human KCNK1 (3ukm^25^) and KCNK4 (3um7^26^), as experimental structures did not exist. PIK3R1 was modeled using PIK3CD (5itd^27^). *In silico* mutagenesis was performed using FoldX version 4.0^28^ to generate initial 3D models of genomic variants.

**Table 1:**
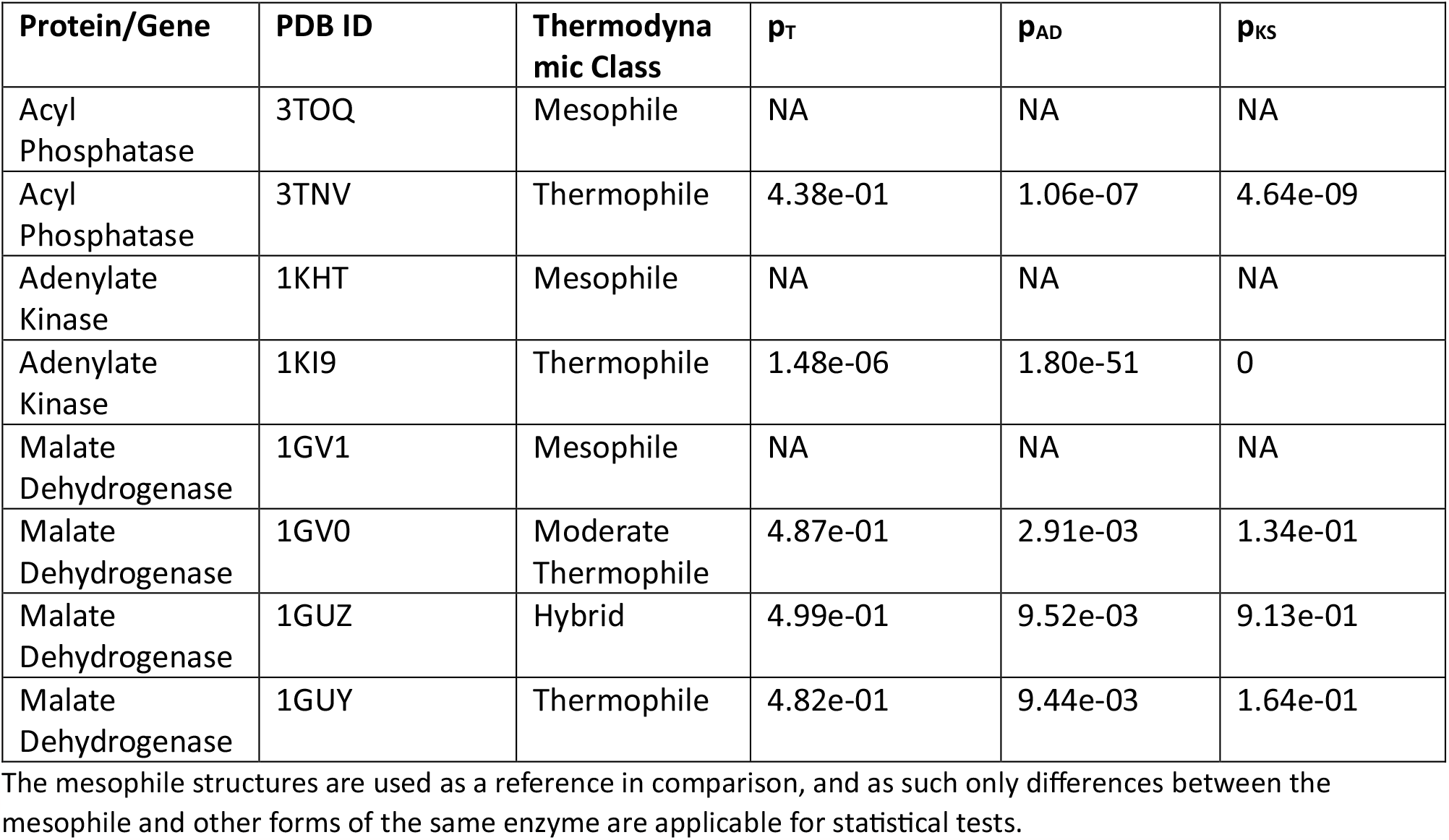
Surface-based comparison of mesophilic and thermophilic enzymes.

**Table 2:** Surface-based comparison of solubilizing variants.

| GENE | Surface Region | PDB ID | Variant | $p_T$ | $p_{AD}$ | $p_{KS}$ |
| --- | --- | --- | --- | --- | --- | --- |
| Granzyme H | Global | 3TK9 | D102N | 7.23e-02 | 6.22e-10 | 8.25e-10 |
| Granzyme H | Local |  | D102N | 3.89e-08 | 1.46e-148 | 0 |
| Granzyme H | Reference | 4GAW | WT | NA | NA | NA |
| IGF1R | Global | 1P4O | E1067A & E1069A | 5.01e-02 | 8.16e-10 | 2.16e-12 |
| IGF1R | Local |  | E1067A & E1069A | 8.71e-34 | 3.55e-299 | 0 |
| IGF1R | Reference | 1M7N | WT | NA | NA | NA |
| UROD | Global |  | D306Y | 3.76e-01 | 7.62e-04 | 6.16e-02 |
| UROD | Local |  | D306Y | 2.32e-02 | 6.28e-19 | 0 |
| UROD | Reference | 3GVQ | WT | NA | NA | NA |

**Table 3:** Local surface-based comparison of missense genetic variants using static structures.

| Protein | Variant | $p_T$ | $p_{AD}$ | $p_{KS}$ |
| --- | --- | --- | --- | --- |
| UROD | D306Y | 2.3E-02 | 6.3E-19 | 0.0E+00 |
|  | E218K | 1.0E-01 | 3.9E-11 | 2.3E-06 |
|  | F232L | 4.8E-01 | 8.7E-01 | 3.3E-02 |
|  | G25E | 1.5E-01 | 5.4E-05 | 2.6E-07 |
|  | G156D | 3.4E-01 | 3.3E-04 | 6.9E-02 |
|  | G318R | 7.0E-03 | 3.4E-23 | 1.1E-16 |
|  | G318R* | 1.5E-02 | 1.3E-22 | 5.4E-08 |
|  | I260T | 5.0E-01 | 3.1E-01 | 5.7E-01 |
|  | K297N | 9.7E-02 | 2.6E-07 | 1.5E-07 |
|  | K297N* | 1.2E-02 | 2.8E-15 | 5.8E-12 |
|  | L253Q | 5.1E-01 | 5.4E-01 | 3.1E-01 |
|  | R144P | 4.3E-01 | 6.5E-01 | 3.3E-02 |
|  | R193P | 4.3E-01 | 6.1E-04 | 3.8E-02 |
| PIK3R1 | E489K | 4.5E-04 | 2.1E-42 | 0.0E+00 |
|  | DQYdel | 2.7E-01 | 3.3E-06 | 6.1E-05 |
|  | N564K | 2.1E-01 | 1.5E-01 | 7.3E-04 |
|  | S393F | 5.0E-01 | 4.2E-01 | 5.0E-01 |
|  | M326I | 5.2E-01 | 8.8E-01 | 2.5E-01 |
|  | MWdel | 4.8E-01 | 5.6E-01 | 6.5E-01 |
|  | F487S | 5.1E-01 | 4.3E-01 | 2.0E-02 |
|  | R409Q | 4.7E-01 | 4.6E-02 | 7.8E-02 |
|  | N564D | 3.0E-01 | 2.4E-03 | 1.1E-06 |
|  | K567E | 4.7E-03 | 1.2E-17 | 0.0E+00 |

\*UROD G318R was crystallized and deposited as 3gw0. PIK3R1 K297N was crystallized and deposited as 3GW3. We compared both between experimental and computational models.

**Table 4:** Local surface-based comparison of missense genetic variants using static structures and dynamics ensembles.

| Protein | Variant | Static or Dynamic | $p_T$ | $p_{AD}$ | $p_{KS}$ |
| --- | --- | --- | --- | --- | --- |
| TBL1XR1 | H348R | Dynamic | 4.7E-03 | 7.2E-19 | 1.7E-14 |
|  |  | Static | 4.0E-01 | 2.3E-05 | 8.4E-03 |
|  | D370N | Dynamic | 9.3E-03 | 3.2E-20 | 5.7E-15 |
|  |  | Static | 3.2E-01 | 1.3E-05 | 4.8E-02 |
|  | T327K | Dynamic | 1.8E-02 | 5.7E-17 | 3.5E-09 |
|  |  | Static | 1.1E-01 | 1.0E-07 | 1.5E-03 |
|  | D370Y | Dynamic | 5.9E-02 | 1.7E-08 | 6.1E-10 |
|  |  | Static | 4.8E-01 | 2.6E-03 | 9.7E-03 |
|  | D328G | Dynamic | 2.9E-01 | 6.0E-02 | 4.0E-03 |
|  |  | Static | 2.9E-01 | 3.9E-05 | 5.5E-02 |
|  | Y446C | Dynamic | 4.2E-01 | 3.0E-01 | 2.0E-01 |
|  |  | Static | 4.8E-01 | 5.1E-01 | 9.1E-01 |
| KCNK9 | G236R | Dynamic | 3.1E-06 | 1.7E-68 | 0.0E+00 |
|  |  | Static | 3.2E-06 | 5.3E-55 | 0.0E+00 |
|  | G236R,<br>A237T | Dynamic | 1.1E-07 | 8.6E-70 | 0.0E+00 |
|  |  | Static | 1.6E-05 | 7.1E-51 | 0.0E+00 |
|  | T199A | Dynamic | 5.1E-01 | 8.3E-10 | 1.4E-04 |
|  |  | Static | 4.9E-01 | 9.1E-01 | 6.5E-01 |
|  | R131S | Dynamic | 1.1E-02 | 5.3E-12 | 2.9E-11 |
|  |  | Static | 1.5E-01 | 3.6E-08 | 1.1E-06 |
|  | A237T | Dynamic | 5.0E-01 | 1.7E-09 | 1.5E-03 |
|  |  | Static | 1.2E-02 | 7.0E-01 | 4.2E-07 |
|  | M159I | Dynamic | 2.1E-01 | 1.6E-04 | 8.7E-04 |
|  |  | Static | 4.8E-01 | 3.2E-01 | 9.1E-01 |
|  | M156V | Dynamic | 2.7E-01 | 2.4E-05 | 4.9E-05 |
|  |  | Static | 4.7E-01 | 8.6E-02 | 6.1E-01 |
|  | R131H | Dynamic | 4.4E-01 | 1.8E-01 | 2.0E-02 |
|  |  | Static | 1.3E-01 | 6.6E-05 | 2.6E-09 |
|  | F135del | Dynamic | 4.7E-01 | 2.6E-03 | 6.2E-02 |
|  |  | Static | 4.7E-01 | 2.4E-01 | 8.9E-01 |
|  | M132R | Dynamic | 4.4E-01 | 2.0E-03 | 4.2E-04 |
|  |  | Static | 2.4E-01 | 8.5E-06 | 8.8E-07 |
|  | R150C | Dynamic | 3.3E-01 | 1.1E-09 | 3.5E-04 |
|  |  | Static | 2.2E-01 | 5.7E-03 | 3.9E-03 |
|  | A237D | Dynamic | 2.6E-01 | 1.9E-04 | 6.9E-07 |
|  |  | Static | 1.3E-01 | 1.0E-05 | 9.2E-08 |
|  | F164C | Dynamic | 5.2E-01 | 8.7E-03 | 3.3E-02 |
|  |  | Static | 4.9E-01 | 3.4E-01 | 7.6E-01 |
|  | M249T | Dynamic | 7.1E-02 | 3.8E-11 | 4.5E-06 |
|  |  | Static | 4.7E-01 | 9.2E-02 | 1.4E-06 |
|  | R131P | Dynamic | 2.6E-01 | 1.8E-05 | 4.2E-04 |
|  |  | Static | 6.0E-02 | 1.8E-11 | 5.8E-12 |
|  | Y138H | Dynamic | 2.1E-01 | 2.8E-04 | 4.0E-13 |
|  |  | Static | 5.4E-01 | 3.1E-02 | 6.9E-01 |

When experimental structures for point mutations existed, we compared them to investigate the use of homology-based methods for interpreting the effect of the same genomic variants.

### Molecular Dynamics Simulation Used to Generate Protein Structure Ensembles

Generalized Born implicit solvent molecular dynamics (MD) simulations were carried out for soluble proteins using NAMD ^29^ and the CHARMM36 ^30^ force field, following a similar procedure as previously described ^31,32^. Models for TBL1XR1 and KCNK9, WT, and each genomic variant were used as input to MD simulations. Each model was used to generate three replicates, and each replicate was independently energy minimized for 10000 steps and heated to 300K over 300ps using a Langevin thermostat. A further 10ns of simulation trajectory was generated, and all trajectories were first aligned to the initial WT conformation using Cα atoms. Frames were extracted from the final ¼^th^ of each simulation as representatives of the dynamic ensemble and used in further analysis.

For the membrane-embedded protein KCNK9, we modeled the explicit environment using the Membrane builder in charmm-gui ^33^ and ran explicit solvent MD simulations in NAMD ^29^ and CHARMM27 forcefield ^34^. We followed a similar procedure while maintaining constant total membrane area and system density by equilibrating in NPT before production simulations in NVT.

### Generating Protein Surface and Local Electrostatic Scores

We used EDTsurf^35^ to generate protein molecular surfaces using the Vertex-Connected Marching Cubes (VCMC) algorithm and the Molecular Surface (MS) parameter. We calculated the electrostatic potential for the structures using APBS^36^ and PDB2PQR^37,38^. The electrostatic distribution data is mapped to the molecular surface from EDTSurf to obtain the surface potential values. These values were used to compute and visualize the differences between the variants and the WT. We used the whole protein surface to compare thermophilic and mesophilic proteins. Structures with variants had their surface character focused on a region surrounding the variant of interest. The local surface selection was used to obtain the tessellations from EDTsurf using any amino acids within a minimum pairwise distance of 4Å (**Figure S1**). We then calculated the electrostatic potential values at this subset of surface points. To assess protein dynamic ensembles, the same procedure was followed for each representative frame (also referred to as a conformer), with pairwise interactions calculated for each representative and the combined values from across time used to compare WT to mutations.

### Statistical Assessment of Altered Surface Potentials

We assessed changes in the electrostatic distribution using a resampling-based approach wherein we randomly selected 100 data points from each dataset, compared using one of three statistical tests, and performed 1000 resampling iterations. A resampling approach is necessary for a robust approach because otherwise any surface change could be made to appear more significant by increasing the number of points used to measure the surface electrostatic potential, or the time density from dynamics. We used three tests - the t-test, Kolmogorov-Smirnov (KS), and Anderson Darling (AD) - chosen because their statistics are based on the difference of means, maximum difference in empirical cumulative distribution, and differences among the tails of the empirical cumulative distribution, respectively. The p-values for the t-test, KS, and AD tests are abbreviated as p_T_, p_KS,_ and p_AD,_ respectively. The p-values obtained from these tests were compared to determine if a small change in distributions, such as for genomic mutations, would be considered significant. We used p<0.01 as a threshold to evaluate our data. An analogous process was used for comparing dynamic ensemble representations of the protein surface, wherein resampling was performed across data from all conformers.

## Results

We applied our approach to scoring protein surface changes to a collection of genomic variants, selected to span positive examples with existing functional validation and disease association and those currently uninterpreted (**Figure 1**). Below, we summarize our findings from assessing 14 total proteins. We demonstrate how global and local changes occur for eight proteins with different physiologic operating temperatures, four proteins with solubilizing mutations (Granzyme H and IGF1R), and four proteins harboring 43 genomic mutations. As our goal is to demonstrate the feasibility of standardizing a protein surface-based approach to scoring the effects of genomic mutations, we chose the four proteins to represent different functions: the complex adaptor protein, TBL1XR1, dimeric potassium ion channel, KCNK9, membrane-bound receptor, PIK3R1, and the soluble enzyme uroporphyrinogen decarboxylase, UROD.

### Thermophilic and Mesophilic Enzymes Demonstrate Clear Surface Differences

The evaluation of surface potential distributions showed that all thermophilic and mesophilic proteins differed significantly (p_AD_ < 0.01) (**Figure 2, Table 1**), supporting the feasibility of our hypothesis that differences in protein surfaces can be evaluated in a standardized way. Compared to its mesophilic ortholog, acyl phosphatase demonstrated a shift towards electropositive (**Figure 2A-C**), p_AD_ of 1.06e-07 (**Table 1**), while adenylate kinase shows a strong electronegative shift (**Figure 2D-F**), with p_AD_ of 1.80e-51 (, **Table 1**). Malate dehydrogenase has four published structures along the mesophilic to thermophilic adaptation spectrum. They show an electronegative peak congruent with the host organism’s temperature adaptation (**Figure 2G-K**). Thus, protein surface properties can be sensitive to temperature adaptation and support our central premise that they could be used to score genomic mutations.

**Figure 2:**
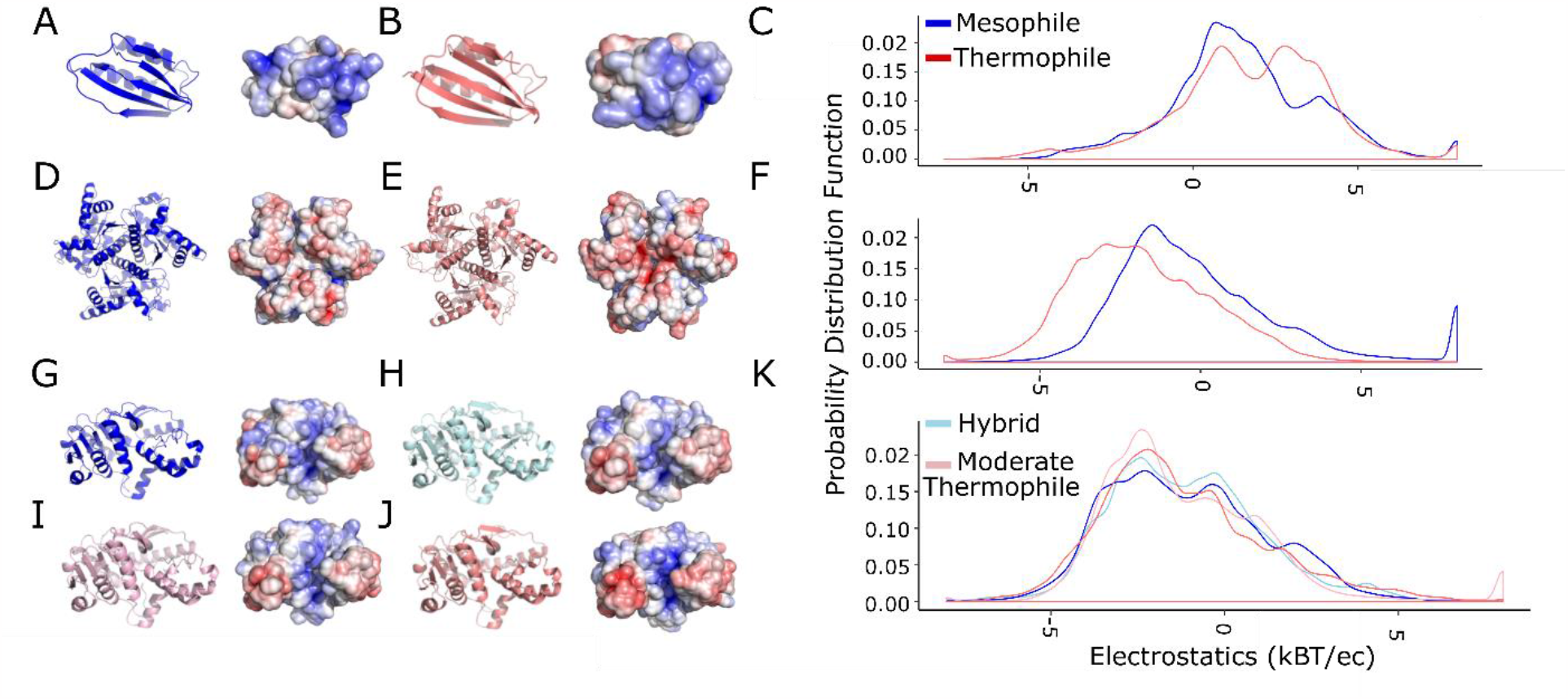
Thermophilic and Mesophilic enzymes demonstrate significant shifts in surface electrostatics. We consider these comparisons as a baseline for defining a considerable change in protein surface electrostatics. We compared 3D models and surface representations for three **A), D), G)** mesophilic enzymes (blue), their **B), E), J)** thermophilic orthologs (salmon), and their **C, F, K)** electrostatic potential probability distributions. These three enzymes are **A-C)**, Acyl Phosphatase, **D-F)** Adenylate Kinase, and **G-K)** Malate Dehydrogenase. Malate dehydrogenase has a hybrid and moderate thermophile structure which shows small changes intermediate between the mesophile and thermophile.

We hypothesized that different statistical tests can be parameterized to reasonably assess the observed differences in protein surfaces. However, even for these positive-control cases, conclusions depended on the type of statistical test used (**Table 1**). For example, t-tests could only distinguish between forms of adenylate kinase but not acyl phosphatase or malate dehydrogenase. The KS test could distinguish among acyl phosphatase forms but not for the other two proteins. Finally, the AD test distinguished among forms of all three proteins. Because the AD and KS tests consider differences in the shape of the surface potential distribution rather than mean differences, they are more able to identify the visually apparent patterns (**Figure 2**). The AD test leverages data from the tails of the empirical probability distribution of each dataset, which in this study is the distribution of electrostatic potential values at the molecular surface and is, therefore, a more robust test for quantifying the patterns that are visually apparent and described above. Thus, comparing mesophilic and thermophilic enzymes indicates that a resampling-based procedure using the AD test is the most promising for evaluating surface potential changes.

### Solubilizing Protein Variants have significant local changes

We next assessed a group of positive control missense variants that have been experimentally characterized and increased protein solubility. They are thus the same type of change that we seek to score, genomic variants, but with a biochemically established effect.

Only minor differences at the mutation site were observed when we assessed the entire protein surface. Therefore, we assessed if the variation was significant at the local electrostatic potential changes caused by the variant. The global surface potential distribution (**Figure 3**) for Granzyme H and IGF1R are statistically significant p_AD_ of 6.22e-10 and 8.16e-10 respectively (**Table 2**). The local surface potential shows the solubilizing variants alter surface properties, in some instances shifting towards a hydrophilic surface in Granzyme H (**Figure 3A-B**) or neutral in IGF1R (**Figure 3D-F**) both of which have an extremely small p_AD_ (Table 2), demonstrating the statistical significance of the protein’s surface charge distribution changes upon mutation. The change in surface potential could explain the improved resolution of crystal structure for Granzyme H from 3Å^39^ to 2.2Å^40^ and IGF1R from 2.7Å^21^ to 1.5Å^41^. All three statistical tests give a significant p-value for local changes caused by the genetically engineered solubilizing mutations. We investigated an additional mutation in UROD and D306Y that has been reported to affect solubility ^19^. The local electrostatic difference in the static WT and D306Y is distinguishable (p_AD_ = 6.28e-19; **Table 2**) and can help us possibly predict the reason for the variant to be insoluble. Therefore, the `-based AD test is also sensitive enough to identify the positive control mutations (Granzyme H and IGF1R) as significant alterations to protein surface properties and helped us evaluate the insoluble genetic mutation in UROD.

**Figure 3:**
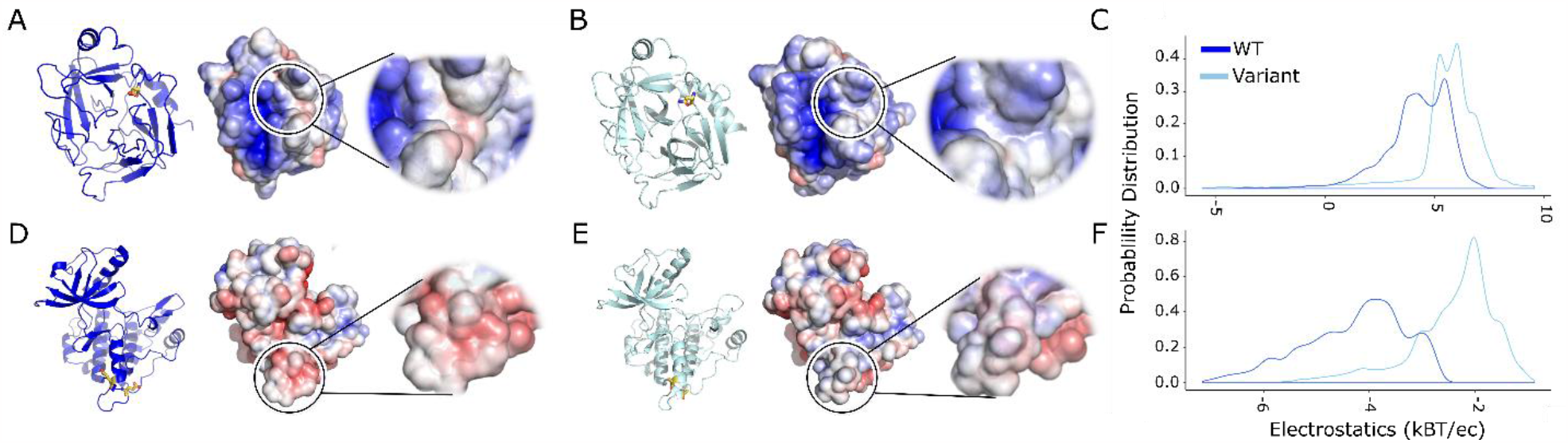
Mutations that increase protein solubility induce significant local changes in electrostatic potential. **A), D)** Wild-type crystal structure with the electrostatic surface shown and zoomed to show the local distribution of charges. **B), E)** 3D structure with the variant shown as sticks in orange followed by the surface shows the change in the local surface compared to the WT. **C), F)** The electrostatic potential distribution shows a significant change between the WT and structure with the variant.

### Dynamic ensembles provide more explicit mechanistic interpretation than static structures

Equipped with the benchmarks above for positive controls, we now assessed the static structure and their corresponding dynamic ensembles (F**igure 1**) for interpreting the effects of missense genetic variation. When assessed globally, static structures show a very small to no difference in the local electrostatic distribution between WT and variants. Of variants, 53% have statistically significant surface charge distribution changes under the AD test for the static structures. UROD (**Figure 8A**) and PIK3R1 (**Figure 8B**) local surface changes showed that when the variant is a charged amino acid or changes from a charged to a neutral amino acid, the potential changes are significant (p_AD_ < 0.01, with a single exception) compared to the WT (**Table 3**). Static structures that capture the nuanced changes observed in the local surface potential. The TBL1XR1 and KCNK9 static median potential change from the median WT were compared to the dynamics median potential change from the median WT (**Figures 4 and 5**). The p_AD_ was again consistently discriminating significant changes, and to a much greater effect when a dynamics ensemble is used (**Table 4**). In the case of KCNK9, average local surface potential shifts for static structures were 0.41 k_B_T/e_c_ with an average p_AD_ of 1×10^−2,^ and the average local surface potential shifts for static structures of TBL1XR1 were 0.43 k_B_T/e_c_ with an average p_AD_ 1×10^−4^. Thus, modeling protein flexibility and the mutation-specific conformational adjustments improved the ability to determine differences statistically.

**Figure 4:**
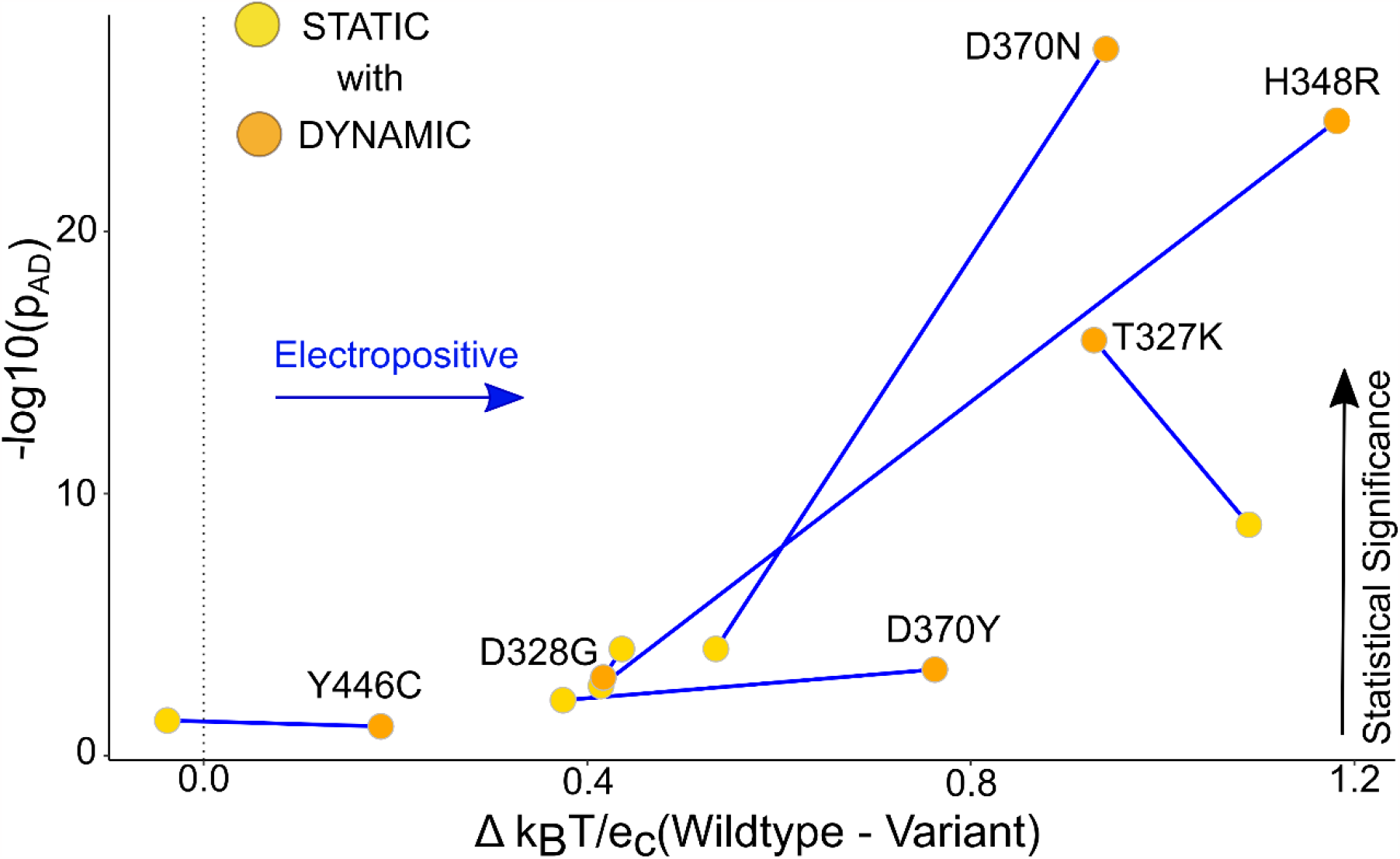
TBL1XR1 Dynamics ensemble helps show a distinguishable change in the surface Potential compared to the Static data. The graph of the median potential difference between the WT and the variant versus the p-value obtained from the AD-test for the static and dynamics ensemble shows that the local surface potential for the ensemble was statistically more significant compared to the static structures. The labels in the graph are assigned to the Dynamic points, and the lines connected show the change in the p-value of Static versus Dynamic.

**Figure 5:**
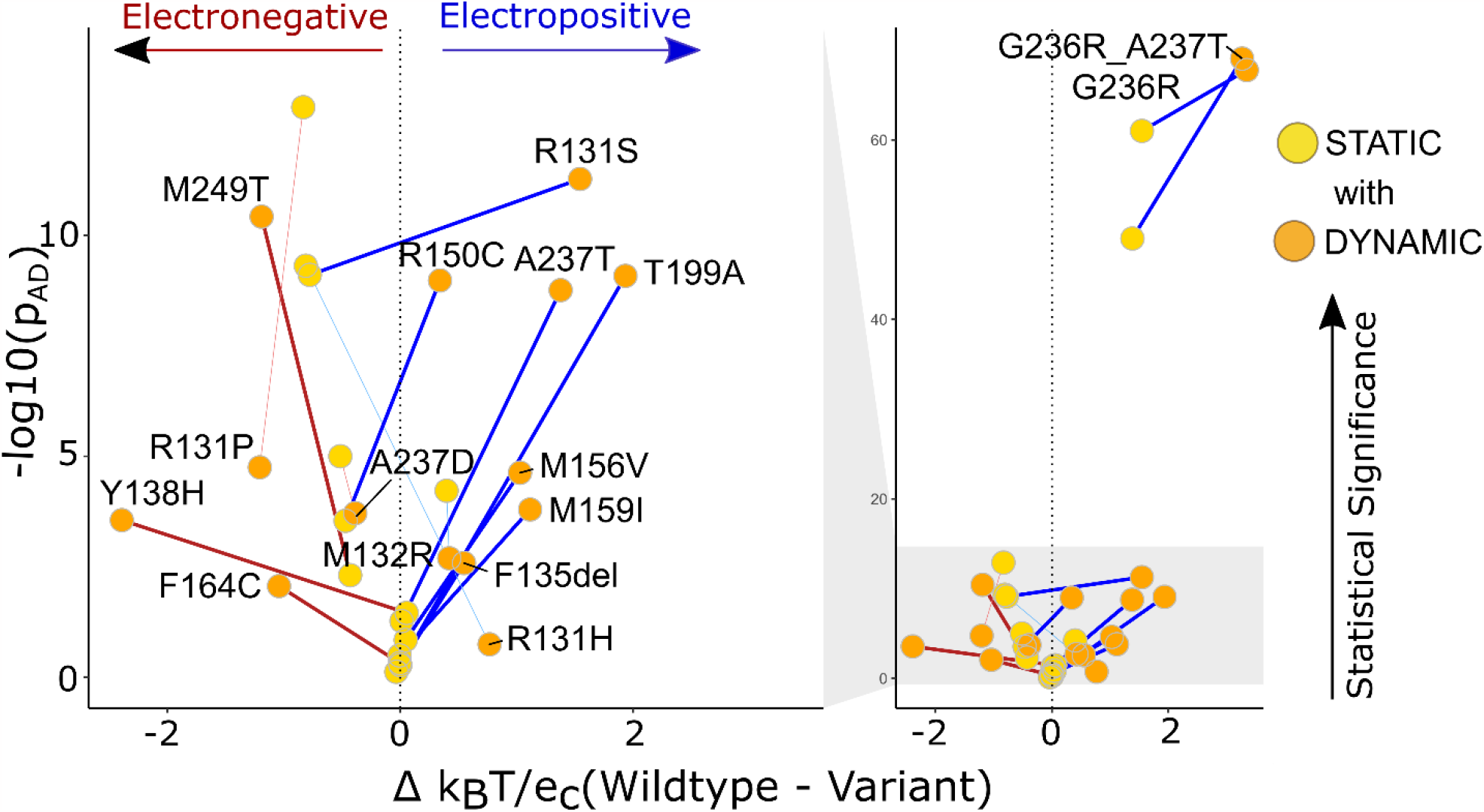
KCNK9 12 variants’ dynamics ensemble showed the local surface potential is better explained by the Dynamics ensemble. The graph represents the distinguishable change in the p-value from low to high (Static to Dynamic) for 12 variants and the 4 variants change from a high p-value to low. The thicker lines connecting the Static and Dynamic variants show the p-value is more significant for the Dynamics ensemble. The thinner lines are the 4 variants that had greater statistical significance in static structures than dynamic ensembles.

Dynamics ensembles help us have a better interpretation of the surface potential data. The local electrostatic potential of the structures obtained from the simulations shows the change compared to the WT in either a positive or negative direction. For TBL1XR1 (**Figure 6B**), we see that 4 out of 6 variants are more Electropositive with a p_AD_ < 0.01, thus possibly causing downstream binding effects for the variants^42^. We also see the difference in surface potential compared to the WT (**Figure 6A**). KCNK9 is a membrane-bound protein channel with a more electronegative surface (WT) for the ions to pass. We see the variants change the surface to either more electronegative or more electropositive (**Figure 7**). The electrostatic surface representation in **Figures 6 and 7** can also be used to compare the differences between the variant and the WT. The representation shows if the surface has moved. In this study, we have only looked at the electrostatic potential; we believe that the protein’s shape also plays a role, especially in protein channels like KCNK9.

**Figure 6:**
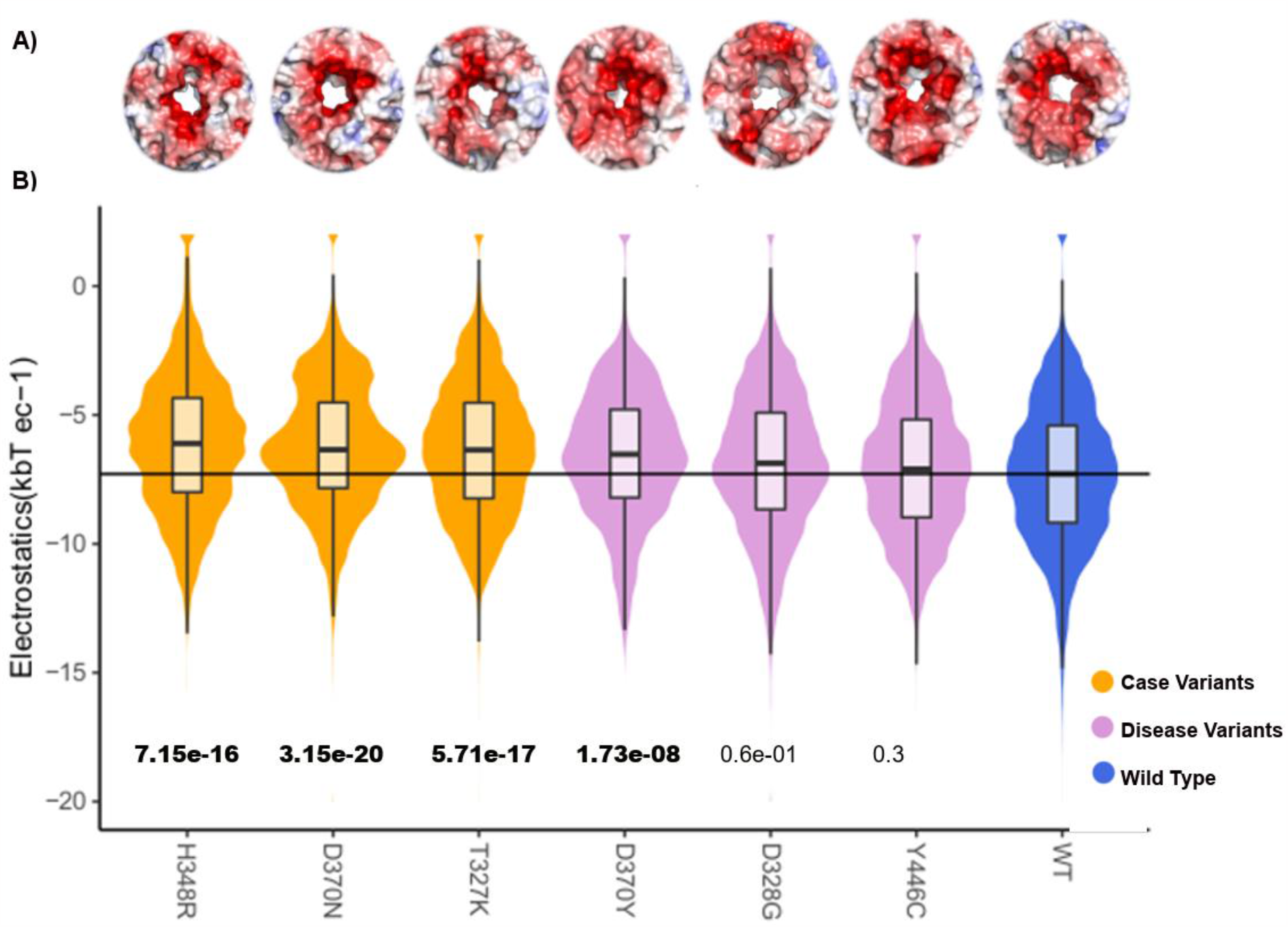
Electrostatic potential for each TBL1XR1 variant dynamics ensemble shows the change in the distribution and median when compared to the WT ensemble. **A)** We show the electrostatic representation of the binding interface of TBL1XR1. **B)** We see the distribution and median of the local surface potential distribution shifts for the 3 case and 1 disease variant as compared to the WT. The variants shift the potential towards the positive direction.

**Figure 7:**
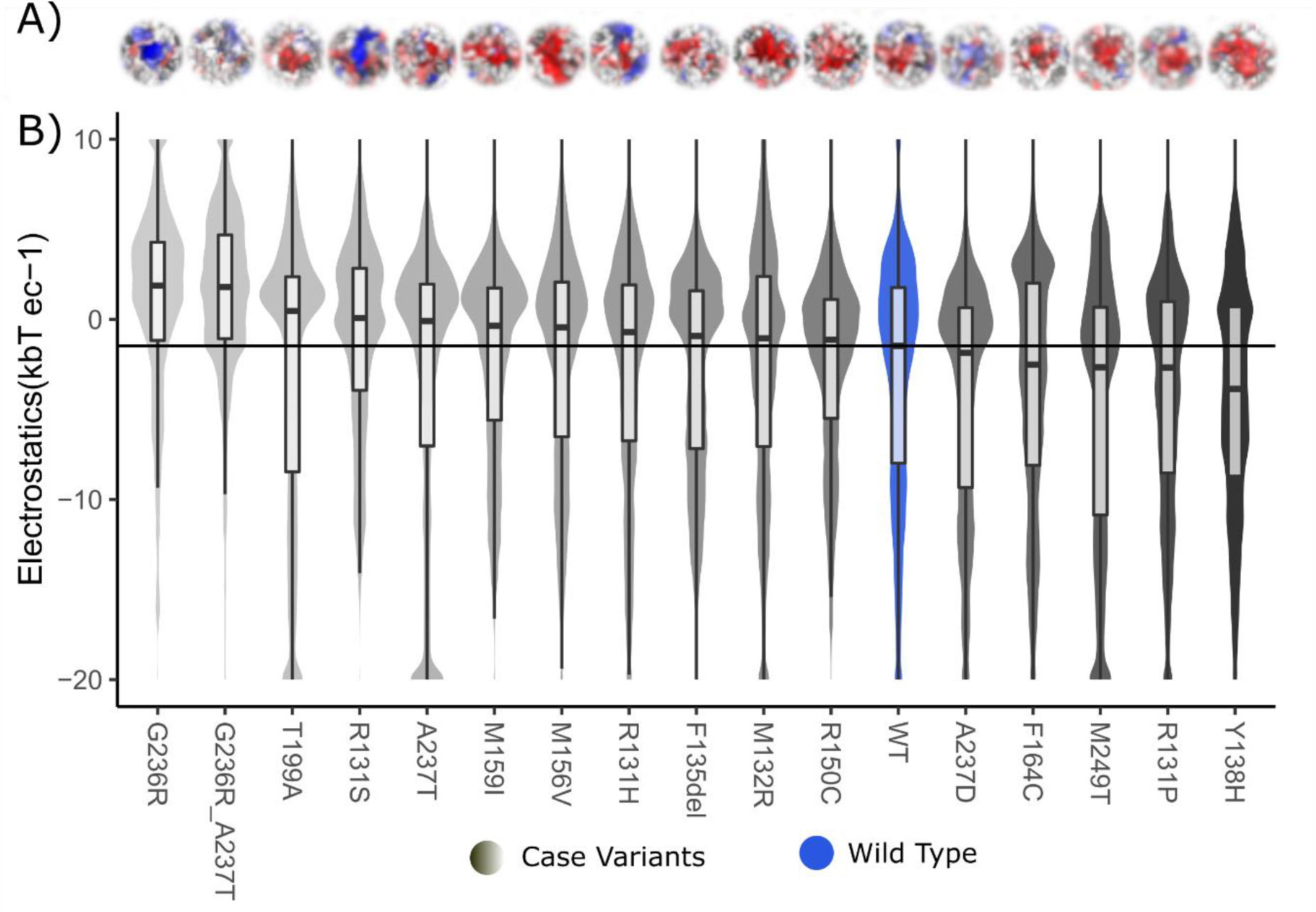
Shift in median surface potential for KCNK9 in both the Electropositive and Electro negative direction as compared to the WT. **A)** We show the electrostatic potential distribution of the pore of KCNK9 membrane protein. **B)** The variants move the potential in both the directions compared to the WT. The color is based on the median values ranging from light grey for more electropositive to dark grey for more electronegative.

**Figure 8:**
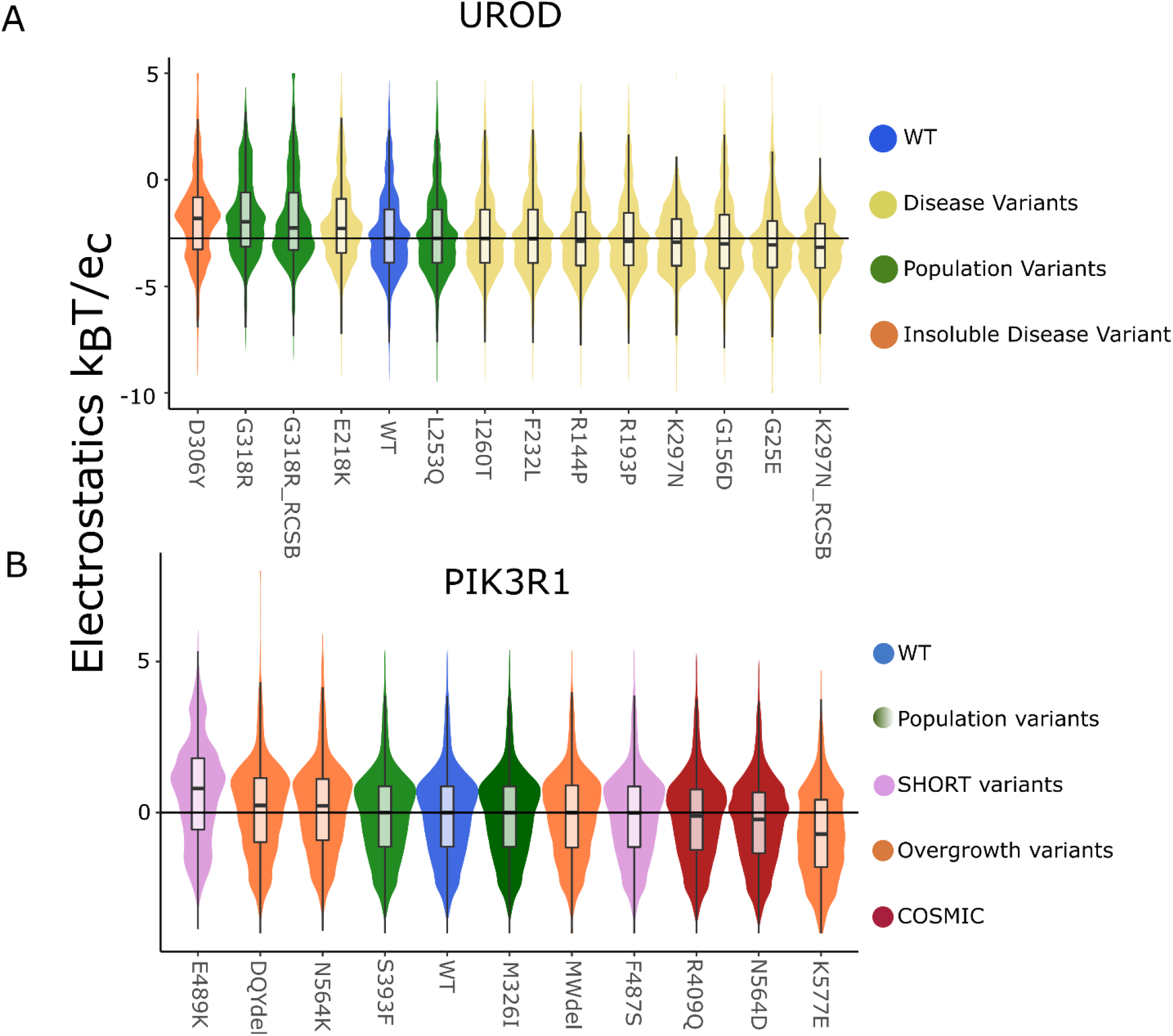
Static local surface potential shows a nuanced change in the variants when compared to the WT. The local surface potential displayed as a violin plot for static structures of UROD and PIK3R1 show small to no difference in the distribution and median with the exception of charged variants.

Scoring dynamic ensembles yielded more information with an average potential shift of 1.15 k_B_T/e_c_ and an average p_AD_ of 1×10^−5^; as for the TBL1XR1 dynamics ensemble, the average local surface potential shift was 0.85 k_B_T/e_c_ with an average p_AD_ of 1×10^−10^. Also, when we look at the statistical significance for TBL1XR1 and KCNK9, we see that 50% of the variants show significant p-values as a static structure. Still, we get about 86% of variants showing significant p-values using dynamics. This helps us show the dynamic ensemble data helps give a better idea about the local surface potential changes in cases where the static structures show a nuanced change.

## Discussion

In this study, we have demonstrated that an objective and standardizable algorithm for assessing meaningful changes in protein electrostatic surface properties can be made. We present our approach, which defines a region of interest – either specific to the mutated site or a particular active site – and monitors how that region of interest is altered directly by the mutation or, with the introduction of dynamic simulation data, indirectly through repacking of the protein fold or allosteric transmission from the mutated site. We identified that dynamics enabled a more straightforward conclusion than static structure alone. Changes to protein surface properties were typically stronger and more statistically significant after accounting for protein dynamics. This type of mechanistic data is not currently predictable from genomic or protein sequences, and we believe it represents a new type of data that can be brought forth for informing the field of genomic data interpretation.

Because protein surfaces are dynamic and influenced by intrinsic and extrinsic factors, it is essential to point out that our goal is to interpret the mechanistic effect of a missense variant, not to identify the quantitative biophysical differences the variant will have in any particular physiologic setting. The latter would require accounting for many extrinsic factors, while we aim for the former, requiring primarily intrinsic factors informed by experimental data. We believe that the process we describe herein can be generalized to the assessment of many types of proteins. We assessed soluble and membrane-bound proteins, characterizing either the effect at the mutated, enzymatic, or functional sites, demonstrating the generalizability of such an approach. Further, surface-based scores can be tailored to account for extrinsic factors that could modulate the effect of a mutated protein in different environments or under other physiologic conditions. Surface-based scores can likely identify significant changes to interfaces and binding sites, without explicitly identifying the molecule(s) that bind. This is an enabling approach for protein science to improve genomic data interpretation.

The data presented in this study indicates an objective assessment of the significance of protein surface changes can be made. Determining an appropriate statistic is important for a new approach. We have shown that standard statistical tests have markedly different performances on protein surface electrostatic potential data. We observed that the change in the peak of the electrostatic potential distribution significantly changes the area under the distribution tails, making the Anderson-Darling test a better indicator of the distribution change compared to the other two statistical tests. We believe that combining permutation-based resampling and the Anderson-Darling test balances sensitivity and false discovery. Thus, we believe changes to protein surface properties could reach standardization befitting use as supportive criteria in the guidelines for interpreting genomic variation. Further, while assessing the surface potential, we found that statistical tests were more sensitive when applied to regions of the protein centered on the altered site rather than when evaluating the entire protein surface. Our future work on protein surface scoring will consider shape-based metrics as well as benchmarking against standardized datasets for protein interactions and broader sets of human disease variation, including from cancer.

## Acknowledgements

This research was completed in part with computational resources and technical support provided by the Research Computing Center at the Medical College of Wisconsin. This publication was supported in part by The Linda T. and John A. Mellowes Endowed Innovation and Discovery Fund and the Genomic Sciences and Precision Medicine Center of Medical College of Wisconsin.

## Supplemental Data

**Supplemental Figure 1:**
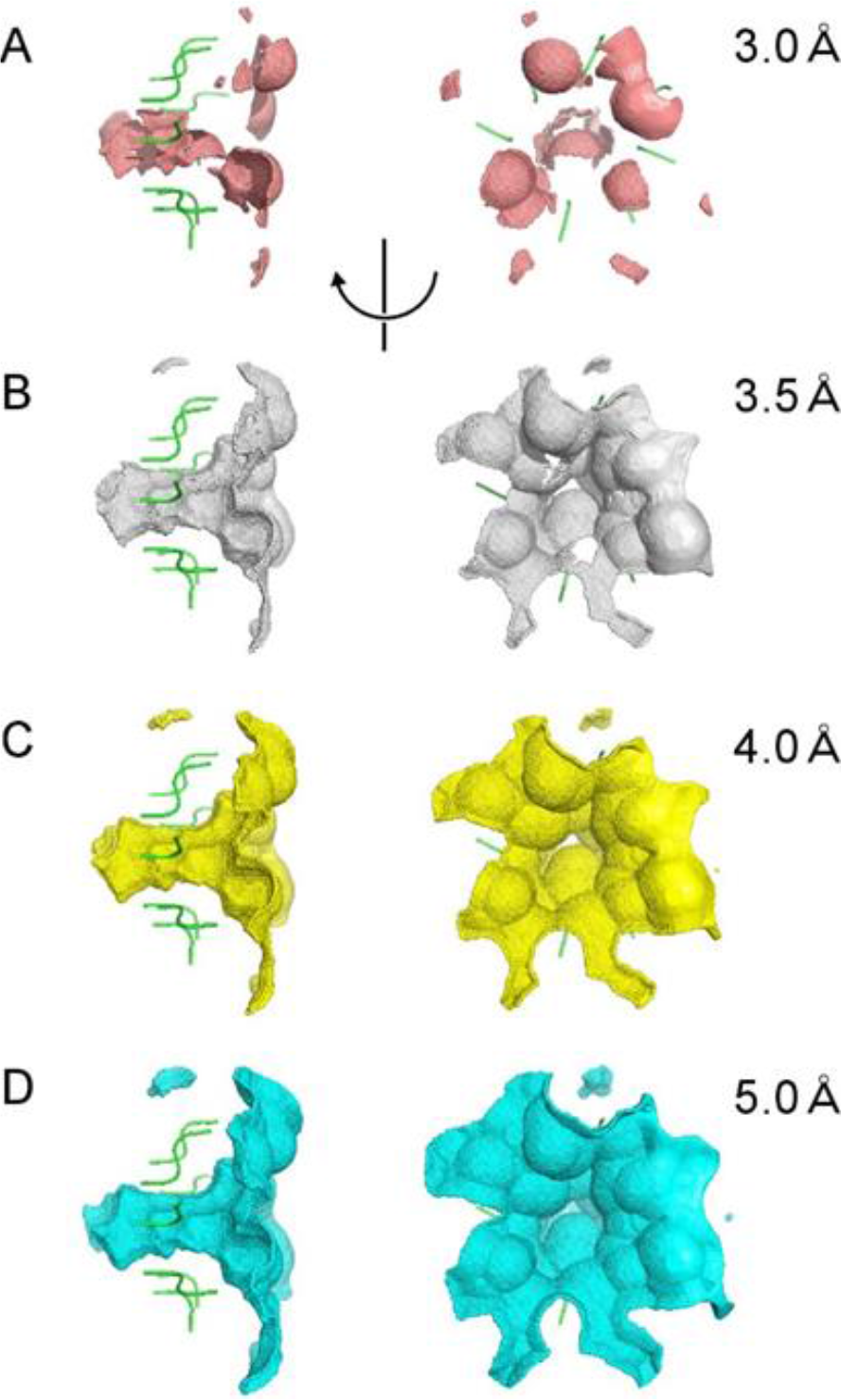
Selection of the subsetted surface and distance. In this study, guided by this example of TBL1XR1, we used 4 Angstrom as a cutoff to select the local surface including the variant and the surrounding environment.

